# Social flocking increases in harsh and challenging environments

**DOI:** 10.1101/2023.08.02.551711

**Authors:** Jessica J. Bellefeuille, Ruchitha C. B. Ratnayake, Emily Cornthwaite, Roslyn Dakin

## Abstract

Grouping with others can provide enhanced information about resources and threats. A key hypothesis in social evolution proposes that individuals can benefit from social information in environments where it is challenging to meet energetic needs. Here, we test this hypothesis by examining the environmental drivers of conspecific flocking behaviour in a large archive of citizen science observations of two common North American birds, the dark-eyed junco (*Junco hyemalis*) and black-capped chickadee (*Poecile atricapillus*). To quantify flocking behaviour, we apply the index of dispersion, *D*, as a metric of clumpiness in each species’ spatiotemporal distribution. We show that juncos in winter are nearly always more clustered than a random expectation, whereas chickadees span a range from uniform to socially clustered distributions. In both species, the degree of social clustering strongly increases with abundance. We identify several key environmental variables that explain the extent of conspecific flocking in both species. Flocks are more socially clustered at higher latitudes, higher elevations, closer to midwinter, and at temperatures that are colder than average given the location and time of year. Together, these findings support the hypothesis that sociality is a key strategy for coping with harsh environments.

**HIGHLIGHTS:**

- Grouping with others can be an important source of information about resources
- We analyzed how flocking behaviour changes throughout winter in two bird species
- We used the index of dispersion to quantify social clustering at a broad scale
- In both species, social clustering increases in response to climate challenges

## Introduction

One of the key benefits of sociality is that conspecifics can provide information regarding food sources and threats (Gager, 2019). Harsh environmental conditions such as extreme temperature and seasonality can increase the challenge of individuals meeting their daily energy budgets (Wingfield et al., 2011). In these harsh conditions, social information may be crucial for locating resources needed for survival (Morand-Ferron et al., 2019). The costs of exploration can be reduced by joining other successful foragers and taking advantage of their resource discoveries (Galef & Giraldeau, 2001). By forming social clusters, individuals can improve their foraging efficiency and fitness (Galef & Giraldeau, 2001; Harpaz & Schneidman, 2020; Jaakkonen et al., 2015). In the evolutionary history of birds, colonial breeding has even facilitated the colonization of harsh environments over long timescales (Cornwallis et al., 2017).

Winter resident birds often experience extreme challenges, including cold temperatures, little to no productivity, and shorter day lengths in which to forage. In these species, a conspecific flock is defined as an aggregation of individuals in the same place at the same time and can arise due to the attraction of individuals to a specific location, resources, or from a mutual attraction between individuals (Emlen, 1952). In birds, increased flocking behaviour has been described during times with cold temperatures, increased snow cover, abnormally low levels of precipitation, inclement wind conditions, and low levels of light (Emlen, 1952; Grubb, 1987; Nakamura & Shindo, 2001). While these observations are consistent with the hypothesis that social grouping is a response to harsh conditions, the previous studies are limited to local populations and narrow geographic ranges.

Here, we aim to test the hypothesis that social flocking is a plastic response to harsh environments by quantifying the dynamics of flocking behaviour at a continent-wide scale (Figure 1A). We focus our study on two winter resident birds that are broadly distributed in North America: the dark-eyed junco (*Junco hyemalis*) and black-capped chickadee (*Poecile atricapillus*). Both species spend spring and summer in pairs or small groups and can form social flocks in winter (Nolan Jr. et al., 2020; Smith & Van Buskirk, 1988). Both species are also extremely common in North America and tolerate a broad range of winter conditions (Figure 1A). To test how these conditions drive conspecific flocking dynamics at a broad scale, we use observations from Project FeederWatch (PFW), a citizen science program that documents birds at backyard bird feeders and other feeding stations throughout winter (Bonter & Greig, 2021). Given the wide geographical distribution of PFW observers and familiarity of these two common species, this program provides an opportunity to examine the ecological drivers of flocking behaviour.

**Figure 1.**
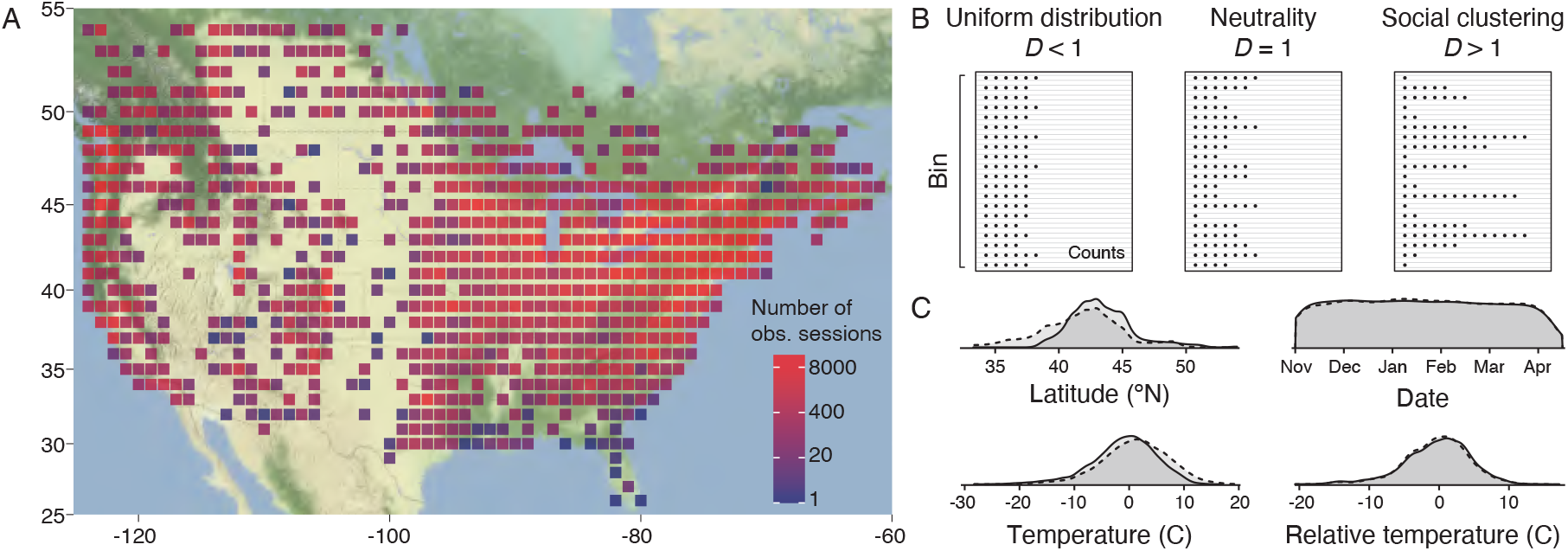
Investigating broad-scale social clustering using two winter resident birds, the dark-eyed junco and black-capped chickadee. (A) The distribution of Project FeederWatch observation sessions for the two focal bird species, binned into 1° latitude x longitude sites. Cell colours indicate the sample size of FeederWatch sessions where one or both focal species were observed. (B) We quantified social clustering of each focal species using the index of dispersion, *D*. Each rectangle in (B) represents a theoretical spatiotemporal bin for a given focal species (i.e., 1° latitude x longitude site over a period of one week). Each row within a bin shows an observation session and the count of individuals that were observed together during that session. When a species is uniformly distributed in space and time, the variance in sample counts is less than the mean count, and *D* < 1. If a species is neutrally distributed, the variance in sample counts scales with the mean (a Poisson distribution), and *D* = 1. When individuals are clustered together in space and time, the variance exceeds the mean, and *D* > 1. (C) Both focal bird species occur over a broad range of winter conditions. In each density plot, the dashed line represents the dark-eyed junco data (n = 9,822 bins), and the solid line represents the black-capped chickadee data (7,851 bins).

We use the index of dispersion, *D*, to quantify the degree of social clustering at a broad scale based on Project FeederWatch observations (Figure 1B). The index of dispersion is a widely used metric of ecological distributions, calculated from a series of counts by dividing the variance of those sample counts by the mean (Perry & Mead, 1979). If birds on the landscape are occurring independently of the presence of others (i.e., social neutrality), *D* will take values centered on one, consistent with a Poisson distribution. If individuals are more clustered together than expected by chance, then the variance in counts will exceed the mean, and *D* will take values greater than one. If individuals are uniformly distributed on a landscape, then the variance in counts will fall below the mean, and *D* will take values less than one. Uniformity can occur if individuals are territorial, or if a species occurs in groups, but the group sizes are highly uniform and/or territorial. Importantly, *D* is a continuous metric that can be compared across a range of scenarios, to evaluate changes in the extent to which a population is clustered.

We apply the index of dispersion to how the winter environment drives social behaviour in dark-eyed juncos and black-capped chickadees. First, we characterize the range of *D* values observed in each species. Second, we examine how environmental factors drive variation in social clustering. We consider four predictors of *D*: latitude, elevation, seasonality, and relative temperature, all of which are expected to influence the ability of small birds to meet their daily energetic needs. Specifically, lower temperatures, higher latitudes, higher elevations, and mid-winter times are all associated with lower resource availability (vegetation and insects) (Fotis et al., 2018; Gillman et al., 2015; Studd et al., 2021). Higher latitudes and higher elevations also have colder climates, which can increase the time and energy required for thermoregulation and foraging (McNamara et al., 2004; Paquet et al., 2016). During winter, a lack of daylight hours presents an additional challenge by limiting foraging time; daylengths are shorter at higher latitudes and mid-winter. Given this background, we predict that social clustering will increase at lower temperatures, higher latitudes, higher elevations, and at mid-winter. We also consider the possibility that these two focal species may differ in their responses to harshness. Black-capped chickadees accumulate food in extensive caches, whereas dark-eyed juncos do not (Brown et al., 2022; Sherry, 1984). Caches are a crucial food source during resource shortages in winter. Conversely, dark-eyed juncos proactively accumulate body fat prior to the onset of winter (Cooper & Swanson, 1994; Rogers et al., 1994). These strategies may drive differences in the costs and benefits of social clustering in response to winter challenges.

## Methods

### Behavioural observations

We analyzed winter observations of dark-eyed juncos and black-capped chickadees collected by volunteers with Project FeederWatch, a project designed to monitor avian winter populations and behaviour. Participants in this project monitor backyard birds at a single location using a standardized protocol and tally sheet (Bonter & Greig, 2021). For each bird species that a participant detects in an observation session, they record a tally of the maximum number of individuals of that species seen simultaneously. In any given case (e.g., 5 black-capped chickadees seen together), it is not known whether the individual birds are persistent flockmates or solitary individuals who happen to be in the same place at the same time (‘gambit of the group’). Importantly, however, by sampling these maximal counts repeatedly across space and time, we are able to measure the extent of clustering in a species’ distribution and how clustering varies through space and time (Figure 1). We expect the extent of clustering to correspond to the use of flocking behaviour by the population. We examined FeederWatch data collected from 2016-2019, with dark-eyed juncos and black-capped chickadees detected in more than 436,000 and 347,000 observation sessions, respectively, during that period.

### Ethical note

All observations of wild birds were collected by citizen scientists while birdwatching as part of Project FeederWatch. No animals were captured or manipulated as part of this study.

### Quantifying social clustering

All data preparation and analyses were performed in R 4.2.1 (R Core Team, 2023). We quantified social clustering separately for each species by grouping species-specific maximal counts into ‘bins’ that covered one week in duration and 1-degree latitude and longitude (representing ∼88-95 km^2^). We did not attempt to infer 0s from sessions where a given species was not detected, because it was important to limit our analysis to conditions that were within the species’ normal winter range and local habitats. Therefore, for each focal species’ analysis, we only used sessions where at least one individual of that species was detected. We computed the index of dispersion, *D*, for each species-bin as the variance in maximal counts divided by the mean. To ensure that estimates of *D* were based on robust sampling, we only used species-bins those that had 10 or more counts of a given species.

To further describe how often *D*-values differed significantly from neutrality, we simulated a large number of neutral (Poisson) samples with varying means, and then removed 0s so that the simulated draws would correspond to the observed data. Then, for each observed species-bin, we drew 100 simulated samples with the same mean and sample size (number of counts). If the observed species-bin had a *D*-value > 95% of the simulated neutral samples, we classified that particular species-bin as significantly clustered. If its *D*-value was < 5% of the simulated neutral samples, we classified it as significantly uniform. If its *D*-value fell within the 5^th^-95^th^ percentile of the expected *D*-values, we classified it as neutral.

### Environmental characteristics

We considered four environmental predictors of social clustering: latitude, elevation, seasonality, and relative temperature. We determined a species-bin’s latitude by rounding to the nearest degree. We determined its elevation by first obtaining the elevation of the coordinates for each observation session within that bin using the R package rgbif, and then taking the species-bin average (Chamberlain et al., 2023). To consider seasonality as a predictor of clustering, we converted each bin’s numerical week to an integer ranging from 1 (beginning in November when winter monitoring commences) to 23 (at the end of April). Hence, the week value captures progress through winter, with mid-winter corresponding to times when day lengths are shortest, and productivity is at a minimum.

We quantified relative temperature as a measure of a bin’s overall weather during that particular week (whereas latitude and elevation above capture the region’s climate, more generally). We first downloaded hourly temperature data archives from the US National Oceanic and Atmospheric Administration (NOAA) database using the package rnoaa in R (Chamberlain & Hocking, 2023). We then determined the daily average temperature for each FeederWatch observation session using its closest NOAA weather station. Most FeederWatch sessions had a weather station within 10-12 km; the small percentage of sessions without a weather station within 35 km were filtered from further analysis. For each species-bin, we then computed the mean temperature in degrees C of FeederWatch sessions in that bin. Finally, we computed relative temperature as (bin mean temperature) – (the mean winter temperatures for that location, defined by its 1-degree latitude and longitude cell). Hence, relative temperature captures how cold or warm it was at that location during a particular week relative to the seasonal average for the location. Negative values of relative temperature represent times that were colder than average, and positive values warmer than average, for a given region.

### Statistical analysis

Because *D* is strongly right-skewed, we used natural log-transformed values for analysis to meet Gaussian assumptions for residuals. Note that when reporting descriptive statistics, we report back-transformed *D* values. To determine which environmental characteristics best explain variation in *D* social clustering, we considered a set of six candidate models outlined in Table 1. Our aim was to identify whether clustering is best predicted by a specific environmental variable, or whether all the full suite of environmental characteristics predict clustering. All candidate models included species as a fixed effect, and the site (categorical) as a random effect. The simplest candidate model had no environmental predictors. The next four candidate models each included a single environmental predictor (either latitude, elevation, seasonality, or relative temperature, respectively). Because mid-winter conditions are expected to be the most challenging, we modelled seasonality (week) nonlinearly as a 2^nd^ degree polynomial. The final candidate model included all four environmental characteristics as joint predictors. To allow for the possibility that the two focal species may differ in their response to these environmental variables, we included interactions between species and each environmental predictor.

**Table 1.** Candidate models to explain variation in social clustering of two bird species.

| Environmental predictors | Model syntax |
| --- | --- |
| None | $\ln(D) \sim \text{species} + (1 \mid \text{site})$ |
| Latitude | $\ln(D) \sim \text{latitude} * \text{species} + (1 \mid \text{site})$ |
| Elevation | $\ln(D) \sim \text{elevation} * \text{species} + (1 \mid \text{site})$ |
| Seasonality | $\ln(D) \sim \text{poly}(\text{seasonality}, 2) * \text{species} + (1 \mid \text{site})$ |
| Rel. temperature | $\ln(D) \sim \text{rel-temp} * \text{species} + (1 \mid \text{site})$ |
| All env. predictors | $\ln(D) \sim \text{latitude} * \text{species} + \text{elevation} * \text{species} + \text{poly}(\text{seasonality}, 2) * \text{species} + \text{rel-temp} * \text{species} + (1 \mid \text{site})$ |

All candidate models were fit as mixed-effects regression models using the package lme4 (Bates et al., 2015). The environmental predictors were mean-centered and standardized to facilitate comparison of effect sizes. We used the MuMIn package in R to compare candidate models using Akaike’s Information Criterion and compute evidence weights and R^2^ values (Bartoń, 2023). All model assumptions were verified. We used the visreg package to generate partial residual plots for the best-supported model (Breheny & Burchett, 2017). The sample size for this analysis was 17,673 species-bins from 196 sites.

## Results

Dark-eyed juncos were nearly always more clustered than the neutral expectation (Figure 2A), with a mean *D*-value of 3.82 (95% CI = 3.76, 3.87; n = 9,822 bins), which is significantly greater than the neutral value of 1 (t-test, p < 0.0001). About 92% of the dark-eyed junco bins were significantly clustered, based on a comparison to simulated neutral samples, and less than 0.1% of the dark-eyed junco bins were classified as significantly uniform (∼8% classified as neutral).

**Figure 2.**
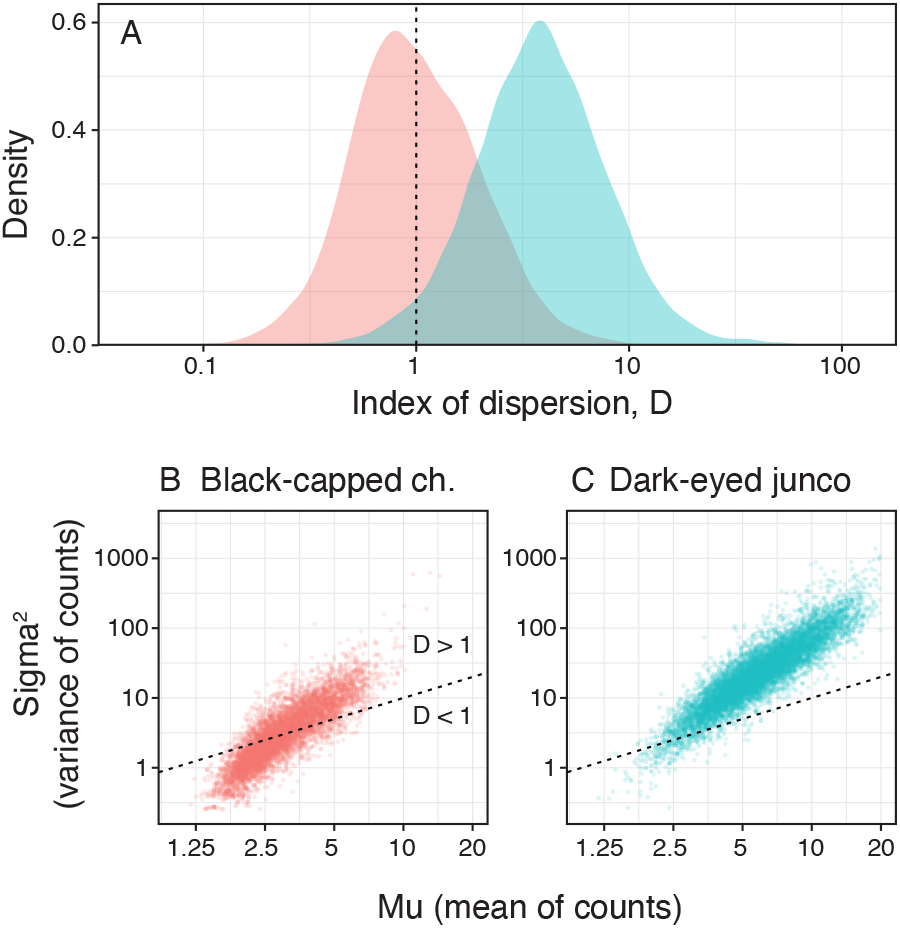
Social clustering in black-capped chickadees and dark-eyed juncos. (A) Distribution of *D*-values for each focal species. Chickadee data are shown in pink (centered on *D* = 1), and junco data are shown in blue (centered on *D* > 1). (B-C) Scatterplots showing the variance in counts (sigma^2^) against the mean count (chickadee n = 7,851 bins; junco n = 9,822 bins). The dashed line in each panel represents neutrality where *D* = 1. These plots reveal that the extent of clustering (i.e., deviation above the neutral line) increases with abundance (mean count) in both species.

In black-capped chickadees, the mean *D* was 0.99 (95% CI = 0.98, 1.01; n = 7,851 bins), which did not significantly differ from neutrality (t-test, p = 0.32). About 38% of black-capped chickadee bins were significantly clustered, and 5% were significantly uniform; the remaining 57% of chickadee bins were neutral. Thus, black-capped chickadees can be found at relatively neutral, clustered, and uniform distributions throughout their winter range (Figure 2A).

In both species, the bins with the greatest mean count also exhibit the largest positive departure from neutrality (as shown in Figure 2B-C), which indicates that clustering in both species increases at times and places when these birds are more abundant.

When comparing candidate models of social clustering, the best-supported model included all four environmental predictors (Akaike weight = 1; Table 2). Notably, the predictors in the best-supported model explain 51% of the total variation in *D* as a measure of social clustering (marginal R^2^_GLMM_).

**Table 2.**
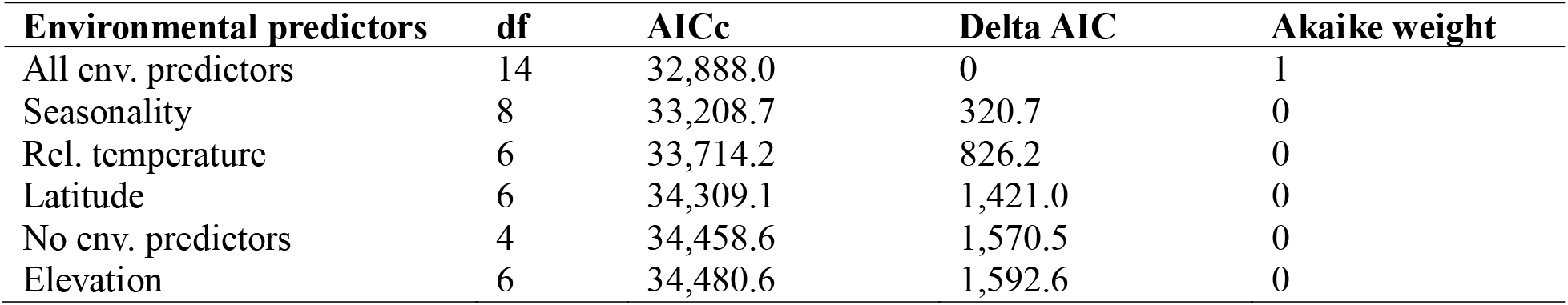
Model selection for social clustering in relation to environmental conditions.

| Environmental predictors | df | AICc | Delta AIC | Akaike weight |
| --- | --- | --- | --- | --- |
| All env. predictors | 14 | 32,888.0 | 0 | 1 |
| Seasonality | 8 | 33,208.7 | 320.7 | 0 |
| Rel. temperature | 6 | 33,714.2 | 826.2 | 0 |
| Latitude | 6 | 34,309.1 | 1,421.0 | 0 |
| No env. predictors | 4 | 34,458.6 | 1,570.5 | 0 |
| Elevation | 6 | 34,480.6 | 1,592.6 | 0 |

The best-supported model is illustrated in Figure 3 and Tables 3 and 4. For both species, social clustering increases at higher latitudes, higher elevations, and during midwinter (Figure 3A-C). The increase in social clustering at higher latitudes is slightly steeper in black-capped chickadees, as compared to juncos, whereas juncos exhibit a slightly steeper increase toward midwinter, as compared to chickadees (Figure 3A, C). Both species also show a significant increase in social clustering when conditions are colder than usual (Figure 3D and Table 4).

**Figure 3.**
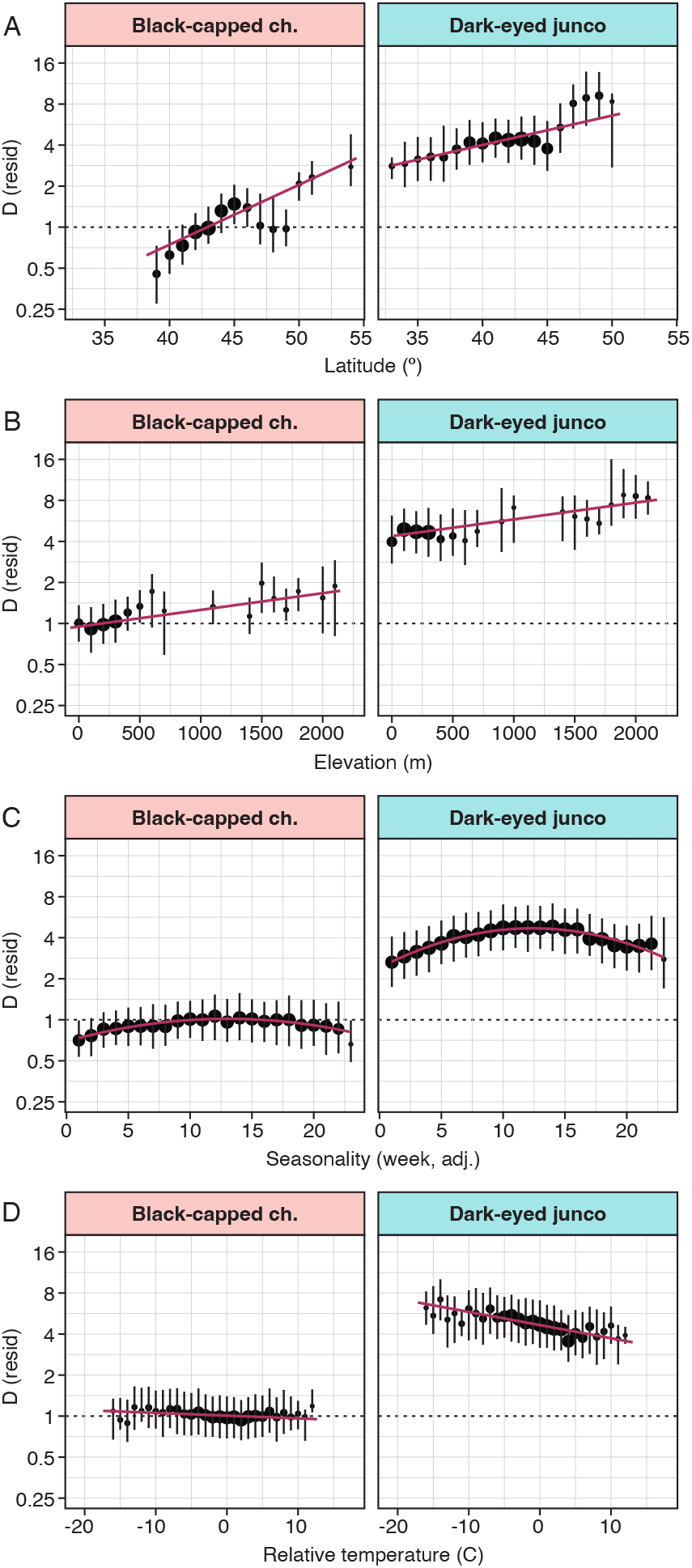
Impact of environmental harshness on social clustering. Each panel shows a partial residual plot for one of the environmental predictors (i.e., adjusting for the other predictors in the best-supported model; n = 17,673 species-bins from 196 sites). For visualization purposes, the x- predictors are grouped into discrete values, with average residual *D*-values shown on the y-axis (error bars are 25^th^ and 75^th^ percentiles). The sizes of the datapoints are proportional to the number of species-bins.

**Table 3.** Best-supported model of social clustering in relation to environmental conditions. Fixed-effect coefficients are provided for the best-supported model of ln-transformed *D*, which included all four environmental predictors and their interactions with species type (n = 17,673 species-bins from 196 sites). All environmental predictors were mean-centered and standardized to facilitate comparison.

| **Fixed effect** | **Estimate** | **95% CI** | **t** | **p-value** |
| --- | --- | --- | --- | --- |
| Species (junco – chickadee) | 1.48 | 1.46, 1.50 | 143.7 | < 0.0001 |
| Latitude STD | 0.30 | 0.25, 0.35 | 12.4 | < 0.0001 |
| Elevation STD | 0.10 | 0.05, 0.15 | 3.4 | < 0.0001 |
| Poly(Seasonality STD-1) | 5.56 | 3.78, 7.34 | 6.1 | < 0.0001 |
| Poly(Seasonality STD-2) | -10.50 | -12.55, -8.46 | -10.06 | < 0.0001 |
| Rel. temperature STD | -0.02 | -0.04, -0.01 | -2.8 | 0.006 |
| Latitude : Species | -0.16 | -0.18, -0.12 | -11.0 | < 0.0001 |
| Elevation : Species | -0.01 | -0.03, 0.01 | -0.9 | 0.37 |
| Poly(Seasonality-1) : Species | 0.58 | -1.83, 2.99 | 0.47 | 0.64 |
| Poly(Seasonality-2) : Species | -9.85 | -12.60, -7.10 | -7.0 | < 0.0001 |
| Rel. temperature : Species | -0.08 | -0.10, -0.06 | -7.7 | < 0.0001 |

**Table 4.** Species-specific slope estimates for social clustering in relation to environmental predictors. These estimates were derived from two separate models fit for black-capped chickadees (n = 7,851 bins from 130 sites) and dark-eyed juncos (n = 9,822 bins from 182 sites). Each predictor had been centered and scaled to a pooled mean of 0 and pooled SD of 1.

| **Fixed effect** | **Black-capped chickadees** |  | **Dark-eyed juncos** |  |
| --- | --- | --- | --- | --- |
|  | **Estimate** | **95% CI** | **Estimate** | **95% CI** |
| Latitude STD | 0.22 | 0.14, 0.31 | 0.15 | 0.09, 0.21 |
| Elevation STD | 0.03 | -0.05, 0.11 | 0.11 | 0.05, 0.17 |
| Poly(Seasonality-1) | 3.73 | 2.69, 4.78 | 4.45 | 3.32, 55.8 |
| Poly(Seasonality-2) | -6.92 | -8.11, -5.72 | -15.09 | -16.37, -13.80 |
| Rel. temperature | -0.02 | -0.04, -0.01 | -0.11 | -0.12, -0.09 |

However, the response of social clustering to relative temperature is much stronger in dark-eyed juncos than it is in black-capped chickadees. While the change in social clustering with relative temperature is statistically significant in black-capped chickadees, the effect size is negligible.

## Discussion

Our analysis show that flocks of dark-eyed juncos and black-capped chickadees become increasingly socially clustered in response to harsh winter conditions. Notably, there is no single environmental predictor that explains the dynamics of social flocking throughout winter. Instead, in both bird species, flocking behaviour varies throughout winter in response to multiple aspects of environmental harshness: flocking increases at higher latitudes, higher elevations, midwinter periods, and at times when the weekly weather is colder than average, for a given location. These results indicate that multiple conditions that constrain a small bird’s energy budget also lead to increased social grouping in flocks, consistent with the hypothesis that social information use is an important strategy to cope with harsh winter challenges.

Dark-eyed juncos are nearly always highly clustered with conspecifics in winter (Figure 2). By contrast, while winter resident black-capped chickadees also form flocks, our analysis shows that the distribution of chickadee flocks on the landscape is frequently neutral and occasionally even uniform. Notably, our findings also indicate in dark-eyed juncos, the degree of clustering behaviour may be more sensitive to winter harshness than that of black-capped chickadees, because the juncos exhibited stronger increases in clustering at mid-winter and at colder relative temperatures (Figure 3). The foraging behaviour of each species may explain these major differences. First, unlike juncos, black-capped chickadees store a portion of their food (seeds or insects) as caches in bark crevices or under dead leaves (Brown et al., 2022; Sherry, 1984). By recovering their caches later in the day, chickadees can supplement daily fat deposition independent of group foraging. Furthermore, in large flocks, group members may be more likely to locate and steal from other individuals’ caches (Pravosudov, 2008). To help avoid pilferage, chickadees fly further away and out of sight before caching when conspecifics are present (Pravosudov et al., 2010). This may explain why chickadees flocks are far more neutrally distributed, and less sensitive to seasonality and temperature.

By contrast, dark-eyed juncos do not cache food, and instead balance energy budgets with body fat accumulated through the winter and stimulated by the onset of winter temperatures and snowfall (Rogers et al., 1994; Swanson, 1991). Juncos also make seasonal movements, as many junco populations perform latitudinal or elevational migrations to overwinter at more southerly or lower elevation regions within North America (Nolan & Ketterson, 1990). Social information, collective predator detection, and the dilution of risk conferred through flocking may be more essential for juncos to survive given the absence of individual caching and the need to move to new locations (Galef & Giraldeau, 2001; Lima et al., 1999), particularly for young individuals migrating for their first time (Németh & Moore, 2007, 2014).

While we chose two focal species that were the most abundant winter residents exhibiting social behaviour, flocking during winter is also observed in many other temperate bird species. Future research can include comparative studies measuring sociality across species and environments. Winter flocks can also consist of more than one species. Investigating social relationships between single and mixed-species flocks can reveal patterns in conspecific vs heterospecific social information use (Jaakkonen et al., 2015; Seppänen et al., 2007).

In this study, we demonstrate the role of harsh environmental conditions in driving social aggregation in two common North American birds, the black-capped chickadee and dark-eyed junco. The broad spatiotemporal range and large sample size of our data indicate a predictable plasticity in social behaviour of wintering birds. We focused here on analyzing flocking behaviour at a broad, landscape scale. While aggregation is a fundamental dimension of sociality, groups also feature complex social relationships between individuals (Strickland et al., 2017). Affiliative interactions such as cooperation can lead to preferential relationships and these pairwise associations can vary in their temporal stability (Dakin et al., 2020). Agonistic interactions can also lead to avoidance between spatially overlapping individuals (Chaine et al., 2011; Strickland et al., 2017). Furthermore, an individual’s ability to acquire information can also be determined by the number and valence of its social relationships (Duboscq et al., 2016). A key next step is to investigate how the dynamics of specific social relationships are influenced by environmental conditions. For example, in harsh environments, the importance of group- living to survival may increase tolerance (or decrease avoidance) between usually avoidant birds.

## DATA ACCESSIBILITY

All data and R scripts are available at: https://figshare.com/s/e6dc79a8cfdf7917d57a

The repository will be made public when the final version of the study is published.

## AUTHOR CONTRIBUTIONS

RD and JB designed the study; RD, JB and RR analyzed the data; all authors wrote and edited the manuscript.

## ACKNOWLEDGEMENTS

We are grateful to the Cornell Lab of Ornithology, Project FeederWatch, and the US NOAA for making data available for this research. We especially thank the many volunteers who have contributed their time and observations to Project FeederWatch.

